# Quantitative Membrane Binding Assays Reveal an Inhibitory Role for the BRAF-Specific Region in CRD and Lipid Interaction

**DOI:** 10.1101/2025.08.03.668294

**Authors:** Vasili Revazishvili, Alexia Morales, Bryn Baxter, Julian Grim, Ani Chakhrakia, Andres Jimenez Salinas, Angelica M. Riestra, Young Kwang Lee

## Abstract

BRAF is a serine/threonine kinase and a central effector of the mitogen-activated protein kinase (MAPK) signaling pathway, frequently mutated in cancer. Its activation is tightly controlled by autoinhibitory mechanisms that regulate membrane recruitment and dimerization. The BRAF-specific region (BSR), located at the N-terminus, is known to promote isoform-preferred RAS binding and facilitate dimerization with kinase suppressor of RAS (KSR), yet its role in regulating lipid interaction has remained unexplored. Here, we identify the BSR as a previously unrecognized inhibitory module that attenuates lipid binding by the cysteine-rich domain (CRD). Using quantitative in vitro reconstitution with supported lipid bilayers and fluorescence microscopy, we demonstrate that the BRAF CRD exhibits high intrinsic affinity for phosphatidylserine-rich membranes, but the inclusion of the BSR markedly reduces the membrane binding. We further demonstrate that the inhibitory function of the BSR correlates with its global electrostatic properties rather than a single defined sequence motif. This inhibitory effect of BSR was corroborated in live cells by quantifying plasma membrane localization of BRAF constructs, including the full-length protein. When canonical autoinhibition of CRD—mediated by sequestration within the 14-3-3 dimer—is disrupted by oncogenic mutation or RAF inhibitor treatment, the BSR assumes a compensatory role in repressing CRD–lipid interaction. This additional regulatory layer provided by the BSR prevents RAS-independent membrane recruitment under both physiological and pathological conditions.

**Broad Impact Statement:** Protein–lipid interactions are a fundamental mechanism for regulating the localization and activity of signaling proteins. This study reveals that the N-terminal segment of BRAF acts as an inhibitory module that suppresses lipid engagement by CRD, particularly when canonical autoinhibition is disrupted by oncogenic mutation or inhibitor treatment. This additional layer of regulation provides new insight into BRAF membrane dynamics and may have implications in therapeutic intervention of dysregulated BRAF signaling.

## Introduction

RAF kinases are serine/threonine protein kinases that function as central effectors of the RAS signaling pathway, regulating key cellular processes including proliferation, differentiation, and survival^1,2^. In mammalian cells, the RAF family comprises three isoforms: ARAF, BRAF, and CRAF. These kinases are activated downstream of RAS by translocating from the cytosol to the plasma membrane, where they initiate the MAPK cascade through phosphorylation of their direct substrates, mitogen-activated protein kinase kinases (MEKs)^2^. Among the three isoforms, BRAF is the most catalytically active and is frequently mutated in a range of human cancers, particularly melanoma, where it functions as a prominent oncogenic driver^3,4^.

RAF kinases are composed of modular domains with distinct regulatory and catalytic functions. The N-terminal regulatory region consists of three main domains: the isoform-specific region, the RAS-binding domain (RBD), and the cysteine-rich domain (CRD)^5^. The C-terminal region contains the kinase domain responsible for downstream MEK phosphorylation. Structural studies using cryo-electron microscopy (cryo-EM) and X-ray crystallography have provided detailed insights into the conformational states of RAF, revealing autoinhibited and active configurations^6–10^. In the inactive state, BRAF adopts a closed, cytosolic conformation maintained by intramolecular interactions between the N-terminal regulatory region and the C-terminal kinase domain (Figure 1a). The autoinhibited state is further stabilized by a 14-3-3 dimer, which binds to two conserved phosphoserine motifs located in the CR2 region (pS365) and the C-terminal tail (pS729). These interactions serve as a molecular handcuff, restraining BRAF in an inactive state. Structural evidence shows that the CRD is sequestered by the 14-3-3 dimer, thereby preventing its engagement with the plasma membrane. The RBD remains accessible for RAS binding and can adopt multiple orientations^9,10^.

**Figure 1.**
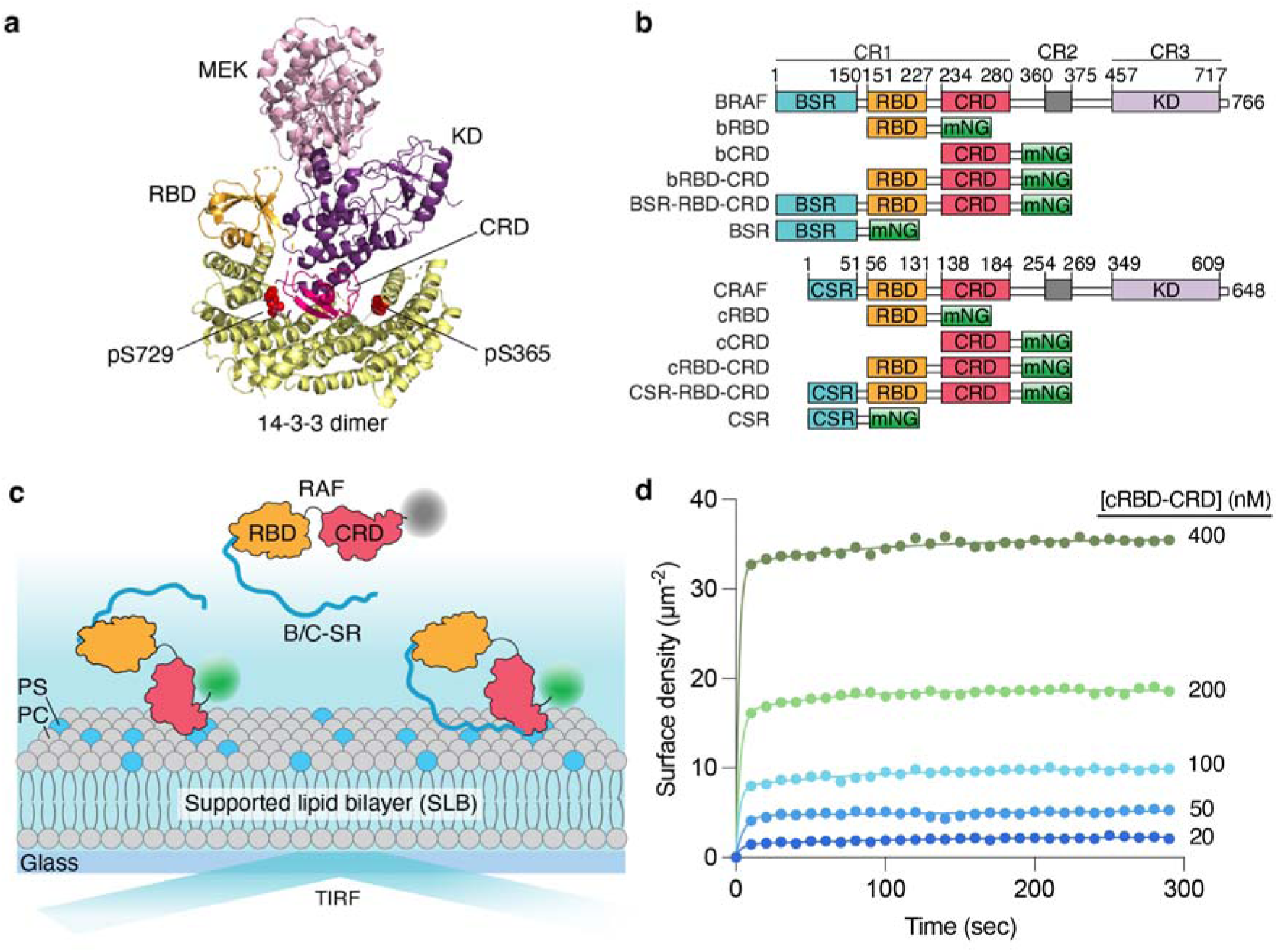
Quantitative characterization of RAF and lipid interactions using SLBs. (a) Structure of the autoinhibited full-length BRAF:MEK1 complex (PDB: 7MFD). The CRD is sequestered within a cradle formed by the 14-3-3 dimer and is not accessible to the membrane. This autoinhibitory architecture is observed across multiple cryo-EM structures with distinct RBD orientations. (PDB: 8DGS, 8DGT) (b) Domain organization of full-length BRAF and CRAF, along with the regulatory domain constructs used in this study. (c) Schematic of the TIRF microscopy–based membrane binding assay using SLBs. (d) Representative membrane-binding kinetics of cRBD-CRD at varying protein concentrations on SLBs containing 80% DOPC and 20% DOPS.

Cellular and biochemical studies have demonstrated that RAF activation is initiated by plasma membrane recruitment^11–14^. RBD–RAS binding and CRD–lipid interaction induce key structural rearrangements that displace 14-3-3, release autoinhibition, and promote RAF dimerization and kinase activation^6,9^. Despite the clear role of CRD in membrane targeting, the regulatory mechanisms controlling CRD accessibility remain incompletely understood.

The N-terminal segment of BRAF, comprising residues 1-150 and hereafter referred to as the BRAF-specific region (BSR), has been implicated in modulating RAF signaling in an isoform specific manner. The selective heterodimerization of BRAF and KSR1 is mediated by direct contact between the N-terminal regulatory region of each protein––the coiled-coiled domain (residues 42-104) within BSR in BRAF and the coiled-coil-sterile α motif (CC-SAM) domain in KSR1^15^. This interaction results in the allosteric activation of BRAF through kinase domain dimerization with KSR1. Additionally, the BSR has been shown to contribute to RAS isoform preference. Bioluminescence resonance energy transfer (BRET) assays in live cells have shown that BSR enables BRAF to preferentially bind KRAS by cooperating with the polybasic region of KRAS^16^. Consistent with these findings, in vitro surface plasmon resonance (SPR) studies have demonstrated high binding affinity between BRAF and KRAS compared to HRAS^17^. Intriguingly, this isoform-preferred interaction requires the presence of the CRD, as deletion of the CRD from the N-terminal region of BRAF abolishes RAS isoform preference^17^. These observations suggest that the BSR and CRD may function cooperatively to regulate BRAF.

Despite its functional importance, the structural and biochemical properties of the BSR remain largely unresolved. Its structure has not been determined in any existing high-resolution structures, either as part of truncated regulatory domains or full-length RAF proteins. Consequently, its contribution to lipid binding and membrane association remains unknown.

In this study, we employed a quantitative in vitro reconstitution system using supported lipid bilayers (SLBs) combined with total internal reflection fluorescence (TIRF) microscopy to characterize the membrane-binding properties of RAF regulatory domain constructs. By systematically varying domain composition and introducing truncations within the BSR, we quantified membrane affinity in terms of surface density and partition coefficients, *K_p_*. Our results demonstrate that the BSR potently suppresses CRD–lipid interaction. The suppressive role of the BSR was further validated in live cells through confocal microscopy, confirming their physiological relevance. Correlative analysis between the electrostatic properties of BSR truncation variants and their membrane-binding affinities revealed that the negatively charged nature of the BSR is a key determinant in suppressing CRD function. Live-cell experiments using full-length BRAF constructs demonstrated that BSR-mediated inhibition and canonical autoinhibition via the 14-3-3 dimer operate in distinct regulatory states of BRAF. BSR-mediated inhibition serves as a compensatory mechanism when canonical autoinhibition is disrupted and BRAF adapts the “CRD-out” preactivating state, as induced by oncogenic mutations or RAF inhibitor binding. Together, our findings identify the BSR as a BRAF-specific autoinhibitory module that suppresses CRD–lipid interactions to prevent RAS-independent membrane recruitment, potentially contributing to regulation under both normal and pathogenic conditions.

## Results

### Quantifying membrane association of RAF on supported lipid membranes

To investigate isoform-specific differences in RAS-independent membrane recruitment, we purified the N-terminal regulatory domains of BRAF and CRAF, excluding their C-terminal kinase domains. Each construct was C-terminally tagged with mNeonGreen (mNG) to enable fluorescence-based detection. We generated a panel of BRAF and CRAF constructs encompassing various combinations of the isoform-specific region, RBD, and CRD (Figure 1b). To assess membrane-binding properties, we employed SLBs containing 20% of the anionic phosphatidylserine (PS) lipids, which is known to play a critical role in recruitment and activation of RAF by interacting with the CRD^18–20^. This two-dimensional membrane system permits quantitative analysis of membrane–protein interaction using TIRF microscopy, which selectively detects membrane-proximal fluorescent species within 100 nm of the membrane surface, effectively distinguishing bound from unbound proteins (Figure 1c)^21^.

To enable direct comparison across constructs, we quantified membrane association by calculating the equilibrium partition constant, *K_p_*. The partition coefficient represents the ratio of protein concentration in the membrane phase to that in the solution phase (see Methods for details)^22^. TIRF microscopy is well suited for determining *K_p_*, as it allows direct measurement of absolute protein surface density when fluorescence intensity is appropriately calibrated^23^. To establish this calibration, we measured TIRF intensity as a function of mNG surface density using fluorescence correlation spectroscopy (FCS). Specifically, His_10_-mNG proteins were tethered to SLBs containing nickel-chelating lipids, and both TIRF and FCS measurements were performed across a range of mNG surface densities (Figure S1). This approach yielded a standard curve showing a linear relationship between TIRF intensity and mNG surface density, enabling us to track membrane-bound protein levels throughout the course of kinetic measurements. All RAF constructs exhibited relatively transient fast membrane-binding kinetics, reaching equilibrium within several seconds. Representative titration biding kinetics are shown in Figure 1d.

### The RBD synergistically enhances CRD-mediated membrane association in BRAF

Travers et al. demonstrated that the tandem arrangement of the RBD and CRD in CRAF synergistically enhances membrane association^24^. To investigate whether BRAF exhibits a similar synergistic property, we measured the partition coefficients of BRAF and CRAF constructs containing the RBD, CRD, or both domains by quantifying their equilibrium surface densities across a range of nanomolar concentrations. As these constructs comprise only the N-terminal RBD and CRD, the observed membrane binding represents the maximal level in the absence of autoinhibitory interactions with the C-terminal kinase domain. The isolated CRAF RBD (cRBD) showed no detectable membrane association at any concentration tested (Figure 2a, square). In contrast, the isolated CRAF CRD (cCRD) exhibited measurable membrane binding, reaching an equilibrium surface density of ∼10 μm^-2^ at 200 nM (Figure 2a, triangle). The tandem CRAF RBD-CRD (cRBD-CRD) construct demonstrated modest enhancement in membrane binding relative to the CRD alone, reaching a surface density ∼20 μm^-2^ at 200 nM (Figure 2a, circle). The enhanced membrane association observed for the CRAF RBD-CRD is consistent with previous SPR measurements^24^.

**Figure 2.**
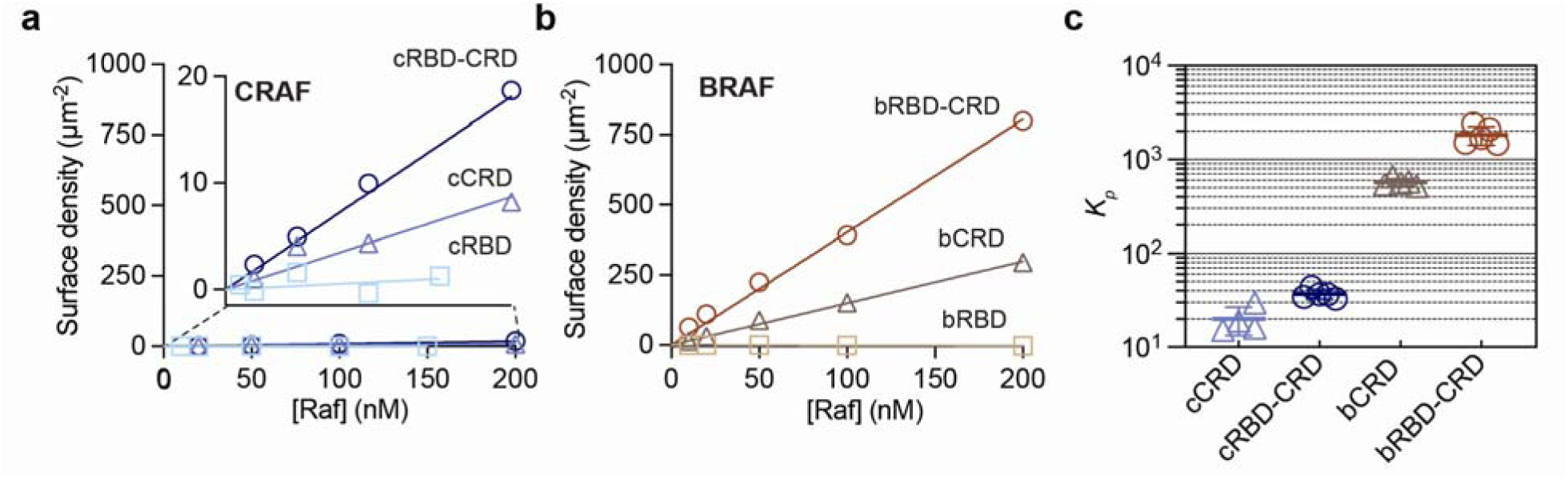
Membrane binding of BRAF and CRAF regulatory domains containing RBD and CRD. Titration binding curves of (a) CRAF and (b) BRAF regulatory domain constructs on supported lipid bilayers composed of 80% DOPC and 20% DOPS. Equilibrium surface densities were obtained from individual binding kinetics and plotted as a function of protein concentration. (c) *K_p_*values of RAF regulatory constructs, calculated from the equilibrium surface density data shown in (a) and (b). *K_p_* values for the isolated RBDs were not determined due to their insufficient membrane-binding affinity. The means and standard deviations were calculated from *K_p_*values determined from individual data points (*n* = 4 or 5) on the titration curves.

The BRAF RBD (bRBD) alone also exhibited no detectable binding to PS-containing membranes similar to CRAF RBD (Figure 2b, square). However, the isolated BRAF CRD (bCRD) showed ∼30-fold higher membrane affinity compared to CRAF CRD, reaching an equilibrium surface density of ∼300 μm^-2^ at 200 nM (Figure 2b, triangle). This marked difference in membrane affinity between BRAF and CRAF CRDs has been reported in previous SPR and live cell studies^25^, supporting the validity of our TIRF-based measurements using supported lipid bilayers. Notably, we observed that tandem BRAF RBD-CRD (bRBD-CRD) achieved an even higher density of ∼800 μm^-2^ (Figure 2b, circle). The results indicate that the RBD of BRAF contributes to enhanced membrane binding of the RBD-CRD tandem, similar to the cooperative effect observed in CRAF. This enhancement may arise from increased local concentrations of anionic lipids near the membrane, facilitated by the RBD when tethered to the CRD, as suggested by molecular dynamics simulations in CRAF^24^. The high membrane association of BRAF CRD and RBD-CRD is reflected in their partition coefficients, determined to be ∼600 and ∼1800, respectively—each approximately an order of magnitude higher than those of the corresponding CRAF constructs (Figure 2c). The relatively lower membrane affinity observed for CRAF is a genuine property of the proteins and not attributable to differences in protein quality. All CRAF constructs were robustly expressed and remained soluble under the experimental conditions (Figure S2). The inherently higher affinity of the BRAF RBD-CRD for PS lipids is likely due to isoform-specific variations in hydrophobic and charged residues, which alter the electrostatic surface potential of the CRD^25^.

### BRAF-specific region suppresses lipid interaction of CRD on supported lipid bilayers

Our quantitative TIRF measurements revealed that BRAF RBD-CRD can achieve a substantially high surface density at physiologically relevant low nanomolar concentrations (<50 nM), solely through lipid interactions and in a RAS-independent manner. The level of membrane binding is comparable to that of CRAF RBD-CRD in the presence of specific interactions with both RAS and PS lipids, measured in a similar supported membrane system^26^. As PS is an abundant lipid in the inner leaflet of the plasma membrane^27^, the high degree of PS-mediated membrane recruitment by BRAF raises the question of how CRD–lipid interactions are regulated to prevent inappropriate activation. One established mechanism involves the formation of an autoinhibited complex comprising BRAF, MEK and a 14-3-3 dimer, in which the CRD is sequestered within a cradle-like structure formed by the 14-3-3 proteins (Figure 1a)^6^. In this conformation, key lipid-interacting residues within the CRD are largely buried by interaction with the 14-3-3 dimer, rendering them inaccessible for membrane binding. Given the more than tenfold higher lipid affinity of BRAF compared to CRAF, we investigated whether an additional regulatory mechanism controlling CRD activity exists specifically in BRAF. Since the isoform-specific region is highly divergent, we tested the role of the BSR in suppressing CRD function. Membrane binding of BRAF and CRAF regulatory domain constructs containing the N-terminal isoform-specific region was quantified on SLBs containing 20% PS lipid. Surprisingly, the membrane affinity of BSR-RBD-CRD in BRAF is approximately twentyfold lower compared to bRBD-CRD, which lacks the entire BSR (Figure 3a). The equilibrium surface density of BSR-RBD-CRD and bRBD-CRD were determined to be ∼40 μm^-2^ and ∼800 μm^-2^ at 200 nM, respectively. This substantial reduction indicates that the BSR negatively regulates the membrane association of the CRD.

**Figure 3.**
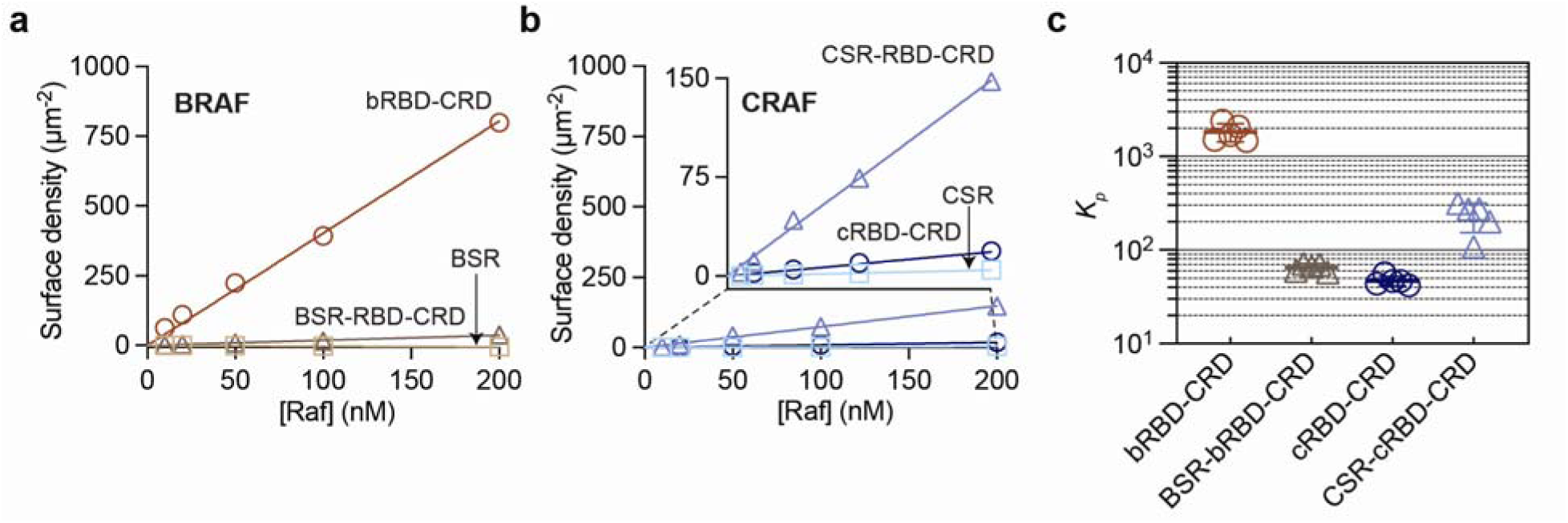
Role of isoform-specific regions of BRAF and CRAF in regulating CRD–lipid interaction. Titration binding curves of regulatory domain constructs from (a) BRAF and (b) CRAF, including constructs with and without the respective isoform-specific regions (BSR and CSR). Membrane binding was measured on supported lipid bilayers composed of 80% DOPC and 20% DOPS. (c) *K_p_* of RAF constructs, calculated from the equilibrium surface density data shown in (a) and (b). The means and standard deviations were calculated from *K_p_* values determined from individual data points (*n* = 4 or 5) on the titration curves.

In contrast, the N-terminal segment of CRAF comprising residues 1-51, referred to as CRAF-specific region (CSR), substantially increases membrane binding (Figure 3b). CSR-RBD-CRD achieved the surface density of ∼150 μm^-2^ at 200 nM, while the cRBD-CRD showed lower lipid binding, reaching ∼20 μm^-2^ under the same conditions. This opposing effect of the BSR and CSR on CRD–lipid interaction may be attributed to their distinct electrostatic properties. The BSR contains a large number of negatively charged residues, resulting in a high net negative charge, whereas the CSR has a near-neutral electrostatic profile (Table S1 and Figure S3). The contrasting effects of BSR and CSR are further reflected in *K_p_* values of BRAF and CRAF constructs, calculated from individual data points in the linear titration binding curves (Figure 3c). Together, these results highlight the differential regulatory roles of the BSR and CSR in modulating CRD–lipid interaction.

### BSR suppresses CRD–lipid interaction in cellular context

To determine whether the isoform-specific differences in membrane binding observed in vitro are preserved in a more complex cellular context, we assessed the membrane localization of mNG-tagged RAF constructs in live HEK293T cells using confocal microscopy. We introduced a mutation in the RBD (R188L for BRAF, R89L for CRAF; denoted with an asterisk) that disrupts its interaction with RAS^25,28^. These mutant constructs allow for evaluation of how the N-terminal isoform-specific region of RAF regulates RAS-independent, lipid-mediated membrane translocation, thereby recapitulating our SLB experiments. For BRAF, the bRBD*-CRD construct lacking the BSR exhibited strong plasma membrane localization, consistent with its high affinity for PS-containing membranes observed in vitro (Figure 4a). In contrast, inclusion of the BSR dramatically reduced membrane localization, with BSR-RBD*-CRD exhibiting a largely diffusive cytoplasmic distribution. For CRAF, both cRBD*-CRD and CSR-RBD*-CRD showed predominantly cytosolic localization, with no appreciable membrane enrichment (Figure 4a). Quantitative analysis of plasma membrane enrichment confirmed that these differences are statistically significant, demonstrating that the BSR suppresses membrane association of BRAF in cellular environments (Figure 4b).

**Figure 4.**
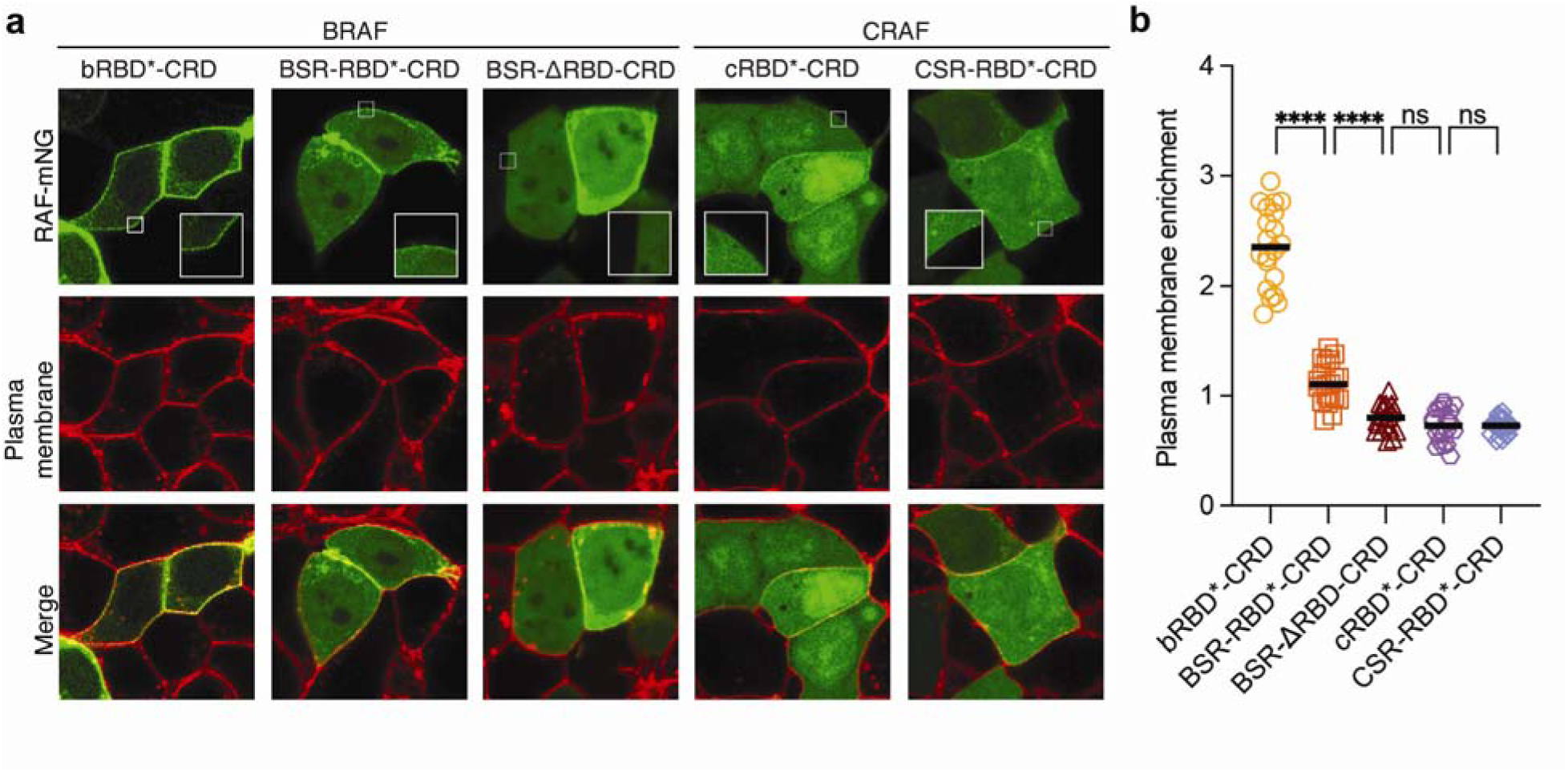
Live-cell imaging confirms the suppressive role of the BSR in CRD–lipid interaction. (a) Representative confocal microscopy images of HEK293T cells transfected with mNG-tagged BRAF and CRAF N-terminal regulatory constructs. All RBDs contain a mutation that disrupts RAS binding (R188L for BRAF, R89L for CRAF). The green channel shows the mNG-tagged RAF proteins and the red channel shows plasma membrane staining with CellMask Deep Red. Insets show magnified views of plasma membranes. (b) Quantification of plasma membrane enrichment for the constructs shown in (a). Enrichment factors were calculated from the average ratio of membrane to cytosolic fluorescence intensity of RAF in individual cells (*n* = 20) Statistical analysis was performed using two-way ANOVA; **** indicates p < 0.0001, ns = not significant.

A previous SPR study reported that the BSR inhibits RBD binding to RAS in an isoform-specific manner^17^. This inhibitory effect requires the presence of the CRD, suggesting functional connection of the BSR with both the RBD and CRD. Because the RBD lies between the BSR and CRD, we tested whether it is required for the BSR to exert its inhibitory effect on the CRD. We examined membrane localization of a deletion construct, BSR-ΔRBD-CRD, in which the entire RBD is removed. BSR-ΔRBD-CRD was predominantly cytosolic with little to no membrane association (Figure 4a,b), confirming that the BSR can directly suppress CRD–lipid interaction independent of the RBD.

### Negatively charged character of BSR globally contributes to suppressive effects on CRD

To identify specific structural features of the BSR that contribute to the downregulation of CRD–lipid interactions, we quantified membrane binding of BSR-RBD-CRD constructs with varying N-terminal truncations. The BSR is largely disordered but contains a structured domain between residues 42–104, which consists of two α-helices that form a dimeric coiled-coil structure. We introduced truncations at three positions—Δ37, Δ105, and Δ127 (Figure 5a). The Δ37 and Δ105 constructs were designed to assess the contribution of the α-helical coiled-coil domain. Hydrogen–deuterium (H–D) exchange experiments have shown that segments of this domain exhibit increased exchange rates upon RAS binding to the RBD (specifically HRAS), suggesting a conformational shift potentially linked to intermolecular interactions^17^.

**Figure 5.**
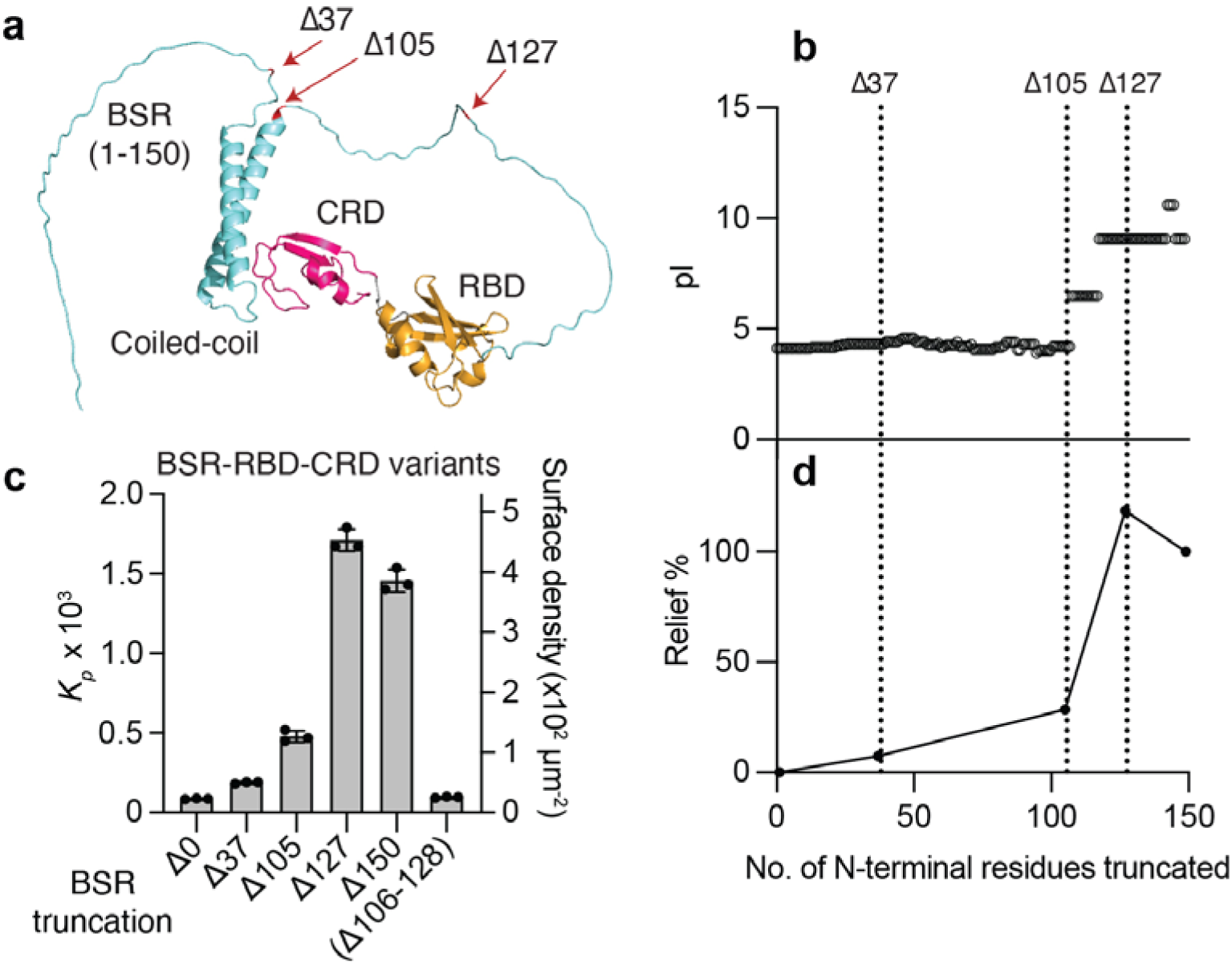
Electrostatic properties of the BSR determine its suppressive effect on CRD–lipid interaction. (a) AlphaFold2 structural prediction of the BRAF BSR-RBD-CRD. Truncation sites within the BSR are indicated with arrows. (b) Isoelectric point (pI) of the BSR as a function of successive N-terminal residue truncations with a single amino acid resolution. (c) *K_p_* values and equilibrium surface densities at 100 nM for BSR-RBD-CRD truncation variants (*n* = 3). SLB composition: 80% DOPC, 20% DOPS. (d) Relief percentage of CRD-lipid inhibition for successively truncated BSR-RBD-CRD constructs, calculated relative to the difference between the full-length BSR-RBD-CRD (Δ0) and ΔBSR-RBD-CRD (Δ150).

The BSR contains multiple negatively charged residues and has an acidic isoelectric point (pI) of 4.13 (Table S1). KRAS-preferred binding of BRAF has been proposed to result from potential electrostatic interactions between the negatively charged BSR and the positively charged hypervariable region (HVR) of KRAS^16^. The negative charge of the BSR may influence CRD–lipid interaction, given that the CRD contains multiple basic residues that mediate membrane association and that the BSR itself may repel the negatively charged plasma membrane. Interestingly, while the pI values of the BSR with gradual N-terminal truncation remain largely unchanged, a sharp increase is observed upon truncation beyond residue 119 (Figure 5b). The Δ127 construct, which retains the basic pI characteristic of the truncated BSR, was generated to test the role of the BSR’s electrostatic properties in modulating CRD–lipid interactions.

To assess the functional impact of these truncations (Δ37, Δ105, and Δ127), we quantified the extent of relief in CRD–lipid interaction. Given that the titration binding curves exhibit a linear relationship between RAF concentration and membrane-bound surface density, the partition coefficient can be accurately determined from single-concentration measurements. *K_p_* values for various truncated constructs were obtained from triplicate measurements at a BRAF concentration of 100 nM (Figure 5c). The relief factor was quantified as the extent to which CRD–lipid interaction was restored upon BSR truncation, calculated using the following equation:

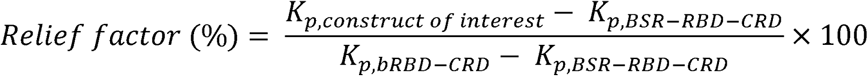

This normalization enables comparison across any construct of interest, including truncation variants, to the full-length BSR-RBD-CRD (Δ0) and the BSR-lacking bRBD-CRD (Δ150) constructs. The Δ37-BSR-RBD-CRD construct, which lacks the disordered N-terminal region preceding the α-helices, exhibited a relief factor of 7% (Figure 5d). Further truncation that removed the α-helical coiled-coil domain in Δ105-BSR-bRBD-CRD modestly restored CRD– lipid interaction, resulting in a 29% relief. Notably, truncation of an additional 22 residues in the Δ127-BSR-RBD-CRD construct led to a relief factor of 118%, indicating complete relief of autoinhibition and enhanced membrane binding compared to bRBD-CRD (Figure 5d). This abrupt shift in relief factor coincides with a transition from acidic to basic pI, along with a corresponding shift in net charge from negative to positive, upon truncation of the first 127 residues (Figure 5b,d and Figure S4), suggesting that the electrostatic properties of the BSR are a key determinant in suppressing CRD–lipid interaction.

Given that the 106–127 region of the BSR, comprising only ∼20 residues, accounts for approximately 70% of the inhibitory effect as observed in Δ105-BSR-RBD-CRD, we generated a targeted deletion variant, Δ(106–127)-BSR-RBD-CRD, to assess the contribution of this specific segment. The targeted deletion Δ(106–127) resulted in only minimal relief (Figure 5c). The disproportionately high contribution of the segment 106–127 observed in the Δ105 construct likely results from spatial proximity to the CRD rather than from indispensable sequence-specific interactions. Since truncation inherently alters steric properties—particularly in intrinsically disordered regions—we cannot fully disentangle electrostatic effects from steric contributions. Nevertheless, the strong correlation between relief factors and pI (or net charge) values across truncated BSR variants supports the conclusion that global electrostatic characteristics of the BSR, rather than a specific segment, play a central role in its inhibitory function.

### BSR assumes a compensatory role in repressing CRD–lipid interaction when canonical autoinhibition is relieved

To determine how BSR-mediated inhibition of the CRD operates in the context of the full-length protein, we first assessed the membrane localization of wild-type full-length BRAF carrying the R188L mutation (FL-BRAF*; an asterisk denotes the mutation in RBD). The R188L mutation in the RBD was introduced to eliminate RAS binding, thereby isolating CRD–lipid interaction from RBD-mediated membrane recruitment. The CRD–lipid interaction was maximally suppressed in full-length BRAF when autoinhibitory inter- and intramolecular interactions were intact. FL-BRAF* exhibited no detectable membrane localization (Figure 6a), with a minimal membrane enrichment factor (Figure 6b, void circle). We next examined the BSR-truncated variant (ΔBSR-BRAF*), which also showed a minimal membrane association (Figure 6a, void square in 6b). These results suggest that CRD sequestration by the 14-3-3 dimer is the dominant mechanism preventing CRD access to the membrane, while the BSR likely plays a supplementary role when the CRD is not structurally accessible.

**Figure 6.**
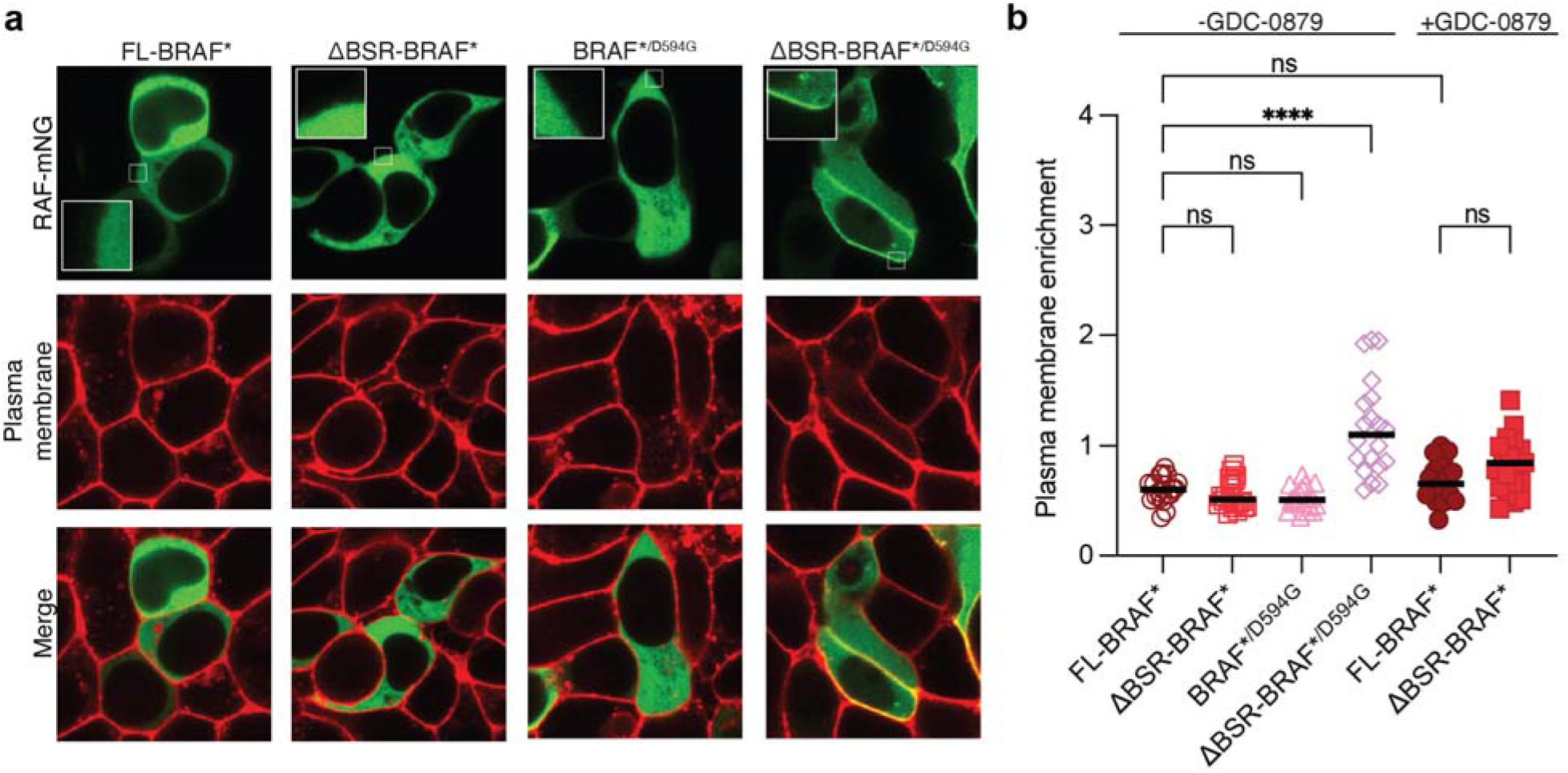
The BSR suppresses CRD–lipid interaction when canonical autoinhibition by 14-3-3 sequestration is disrupted. (a) Representative confocal microscopy images of HEK293T cells transfected with mNG-tagged full-length (FL) and BSR-lacking BRAF (ΔBSR) constructs. Cells expressing wild-type and kinase-inactive oncogenic mutant (D594G) BRAF variants were shown. The RBD contains a mutation (R188L) that disrupts RAS binding. The green channel shows mNG-tagged BRAF; the red channel shows CellMask Deep Red staining of the plasma membrane. Insets display zoomed-in views of plasma membranes. (b) Quantification of plasma membrane enrichment across conditions with or without GDC-0879 treatment. Representative confocal microscopy images for inhibitor-treated cells are presented in Figure S5. Enrichment values were calculated from the ratio of membrane to cytosolic fluorescence in individual cells (*n*=20). Statistical analysis was performed using two-way ANOVA; **** indicates p < 0.0001, ns = not significant.

Based on the observations above, we hypothesize that the BSR suppresses CRD–lipid interaction when canonical autoinhibition is released (i.e. upon liberation of CRD from the cradle formed by the 14-3-3 dimer). To test this hypothesis, we examined a kinase-inactive BRAF mutant D594G (BRAF*^/D594G^) and its BSR-truncated counterpart (ΔBSR-BRAF*^/D594G^). The D594G mutation is a well-characterized kinase-dead class III BRAF mutation found in a broad range of tumor types^29,30^. It resides in the DFG motif of the kinase domain, rendering BRAF catalytically inert and simultaneously disrupting the autoinhibited conformation by displacing the orientation of the kinase domain helix αC in an αC-in (active-like) conformation^31^. This conformational shift leads to a preactivated state (CRD-out), in which the CRD is released and no longer sequestered by the 14-3-3 dimer, as observed in a cryo-EM structure of BRAF ^D594G^ ^32^. Full-length BRAF*^/D594G^ with an intact BSR showed no appreciable membrane localization, despite adapting the CRD-out state (Figure 6a, void triangle in 6b). In contrast, ΔBSR-BRAF*^/D594G^ resulted in a clear membrane localization, confirming that the CRD is capable of engaging with lipids when the BSR is absent (Figure 6a, void diamond in 6b). This difference demonstrates that the BSR alone is sufficient to suppress CRD–lipid engagement. The BSR assumes a compensatory role in inhibiting the CRD upon relief of canonical autoinhibition by 14-3-3. This mechanism provides an additional, previously unrecognized layer of autoinhibition in BRAF.

BRAF inhibitors weaken the CRD-mediated autoinhibitory contact between the kinase domain and the N-terminal regulatory region to varying degrees^33,34^. This reduction in autoinhibition can enhance RAS–RAF interactions and promote RAF dimerization, leading to paradoxical activation of the pathway under certain conditions^35,36^. It occurs through a shared mechanism with oncogenic BRAF activation described above: an inward conformational shift of helix αC induces the CRD-out state^8,32^. To test whether the BSR contributes to suppression of CRD–lipid engagement when autoinhibition is relieved by inhibitor binding, we treated cells with GDC-0879, a type I inhibitor that stabilizes the helix αC-in conformation and is known to induce the CRD-out state^32^. Cells were incubated with 2 μM GDC-0879, and membrane localization of full-length FL-BRAF* and ΔBSR-BRAF* was quantified. Although a statistically significant difference was not detected, we did observe a subpopulation of cells that exhibited membrane translocation of ΔBSR-BRAF*, whereas the entire population of FL-BRAF* cells remained strictly cytosolic (Figure S5, solid symbols in Figure 6b). This observation indicates that the BSR can suppress CRD–lipid interaction when CRD autoinhibition is weakened by RAF inhibitor treatment.

Together, these results highlight dual mechanisms of CRD regulation in BRAF: one mediated by the canonical autoinhibited complex with 14-3-3 and MEK, and a second, BRAF-specific mechanism conferred by the BSR that suppresses CRD–lipid interaction when the CRD is structurally accessible.

## Discussion

Recent studies have established that the BSR plays multifunctional roles in regulating RAF activity. It enables KRAS-preferred interaction of BRAF and facilitates kinase domain dimerization with KSR1 through binding to its CC-SAM domain^15,16^. It also strengthens intramolecular autoinhibition contact between the N-terminal regulatory region and the C-terminal kinase domain^17^.

Using a combination of in vitro reconstitution on supported lipid bilayers, mutagenesis, domain truncations, and live cell imaging, we systematically investigated how the BSR modulates membrane binding. Our study uncovered a previously unrecognized inhibitory function of the BSR: suppression of CRD–lipid interaction. Quantitative TIRF microscopy and partition coefficient measurements revealed that the BRAF CRD has significantly higher membrane affinity than its CRAF counterpart. Importantly, we found that N-terminal segments of BRAF and CRAF have opposite effects on membrane binding. The BSR reduces CRD–lipid interaction whereas CSR enhances it. The BSR-mediated suppression was further validated in live cells, where the BRAF regulatory region exhibited reduced plasma membrane localization in the presence of the BSR. Further analysis through domain truncation and electrostatic profiling revealed that the inhibitory effect of BSR correlates strongly with its net negative charge. Notably, a sharp loss of inhibitory function was observed upon truncation of residues 1–127, coinciding with a shift in pI from acidic to basic, and a corresponding transition in net charge from negative to positive. Neither truncation nor targeted deletion identified a narrow segment responsible for the inhibition of CRD–lipid interaction, supporting a model in which distributed electrostatic features, rather than a single discrete motif, mediate CRD suppression.

Our live-cell study on the BSR in the context of full-length BRAF suggests that its inhibitory function becomes particularly important when the CRD is structurally liberated from the 14-3-3 dimer and accessible to the membrane (i.e. the CRD-out state)—a state thought to occur during the transition from an inactive monomer to an active dimer (Figure 7). This preactivated CRD-out state can be populated under various conditions and is closely associated with pathogenic mechanisms. A recent cryo-EM study^32^ has shown that the CRD-out state is common to all three classes (I, II, and III) of BRAF mutants, including V600E and D594G—the latter of which was tested in the present study. This mechanism is also implicated in the action of RAF inhibitors, which induce a similar preactive-like conformation that facilitates paradoxical activation through dimer formation. We observed that the BSR continues to suppress CRD–lipid interactions to a similar extent as the canonical autoinhibitory mechanism mediated by 14-3-3 dimers, under conditions of oncogenic mutation or inhibitor binding. This provides a mechanism by which BRAF membrane recruitment remains dependent on active RAS binding.

**Figure 7.**
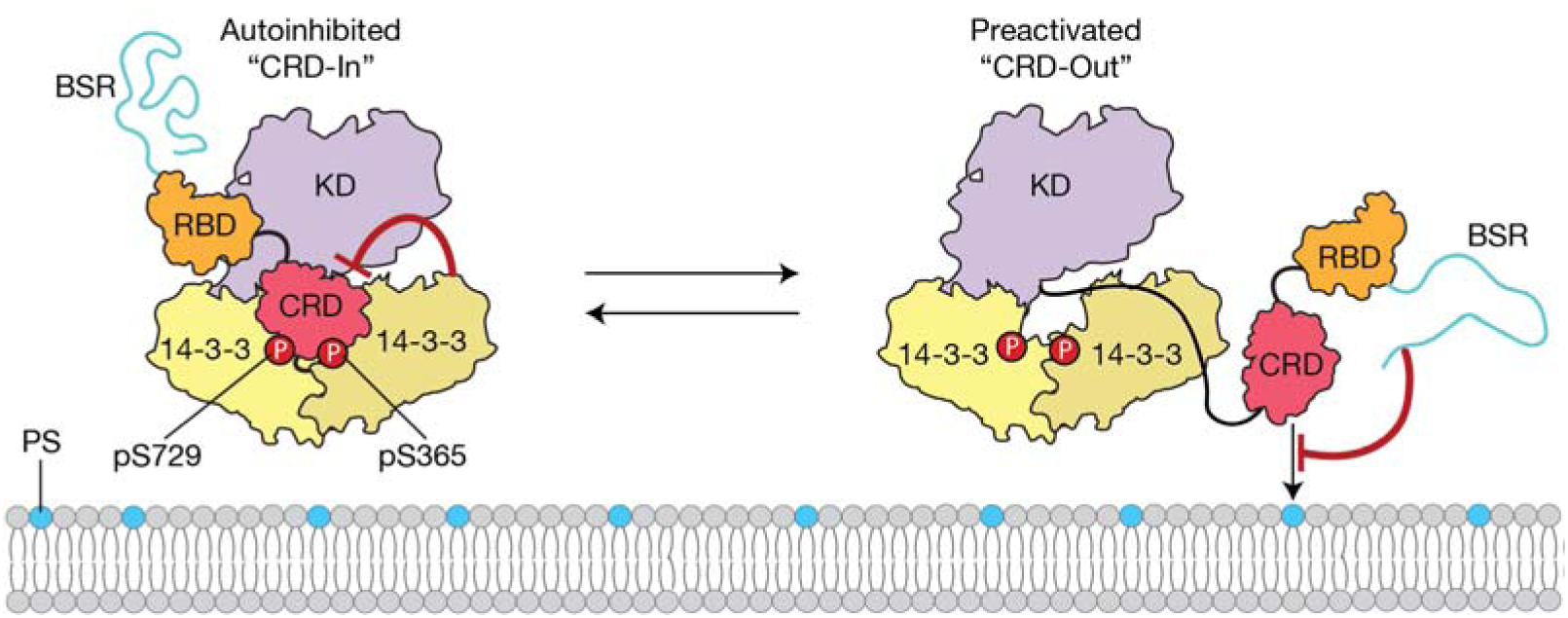
Model illustrating dual inhibitory mechanisms regulating CRD–lipid interaction in BRAF. In the autoinhibited state, CRD–lipid interaction is blocked by canonical autoinhibition, where the CRD is sequestered within a cradle formed by the 14-3-3 dimer, making its lipid-binding residues inaccessible. Upon disruption of canonical autoinhibition—such as through oncogenic mutation or RAF inhibitor binding—BRAF adopts a preactivated “CRD-out” conformation in which the CRD is structurally released. In this state, the BSR continues to suppress CRD–lipid engagement. The electrostatic properties of the BSR have been identified as a key determinant of this suppressive function. PS indicates phosphatidylserine lipids.

The precise mechanism by which the BSR suppresses CRD–lipid interaction remains to be elucidated. One possibility is that the negatively charged BSR engages in intramolecular interactions with the positively charged CRD, sterically or electrostatically occluding membrane-binding residues. Alternatively, electrostatic repulsion between the acidic BSR and the negatively charged plasma membrane may destabilize CRD–lipid association. These two mechanisms may not be mutually exclusive and may operate in concert. Further studies are needed to determine how the inhibitory function of the BSR is relieved under activating conditions. Interactions with positively charged partners—such as the polybasic HVR of KRAS or CC-SAM of KSR1—could potentially neutralize the inhibitory electrostatic effect of the BSR and facilitate CRD engagement with the membrane. Investigating the role of the BSR in the context of RAS isoforms, particularly KRAS, and its impact on membrane-binding kinetics will provide deeper insight into BRAF activation within physiologically relevant membrane environments.

## Conclusions

In summary, we propose that the BSR functions as a multifunctional regulatory module with dual roles in both promoting and constraining BRAF activity. While it facilitates activation through isoform-specific RAS recognition and KSR-dependent dimerization, as demonstrated in previous studies, it also acts as a unique BRAF-specific autoinhibitory element that restricts CRD–lipid interaction. Upon relief of canonical autoinhibition—such as that induced by oncogenic mutations or RAF inhibitor binding—the BSR assumes a compensatory role in suppressing CRD activity. This functional duality adds an additional regulatory layer to the current model of RAF signaling and underscores the potential relevance of the BSR in modulating dysregulated BRAF activity in disease contexts.

## Materials and Methods

### Bacterial expression plasmids

Bacterial expression constructs for CRAF included the following segments: CSR (residues 1-51), cRBD (52-131), cCRD (138-188), cRBD-CRD (52-188), and CSR-RBD-CRD (1-188). For BRAF, constructs included: BSR (1-151), bRBD (151–227), bCRD (228-284), bRBD-CRD (151-284), BSR-RBD-CRD (1-284), and a series of N-terminal truncation variants of BSR-RBD-CRD. cDNA encoding the protein of interest (POI) was cloned into a modified version of the 2C-T vector (Addgene plasmid #29706). In this version, the N-terminal His_6_ tag was replaced with a TwinStrepTagII, and the vector was renamed 2C-T-TwinStrepTagII-MBP-N10-TEV-POI-GGGGS_3_-mNG. The POI fused with C-terminal mNG via a flexible triple GGGGS linker was inserted at the C-terminal TEV cleavage site. All cloning was performed using Gibson Assembly. All open reading frames of bacterial expression plasmids were sequenced by DNA sequencing.

### Mammalian expression plasmids

Mammalian expression CRAF constructs included CSR-RBD*-CRD (aa 1-194) and cRBD*-CRD (aa 52-194). BRAF constructs included bRBD*-CRD (aa 151-290), BSR-RBD*-CRD (aa 1-290), BSR-ΔRBD-CRD (aa 1-290 with deletion of 155-227), FL-BRAF* (aa 1-766), and ΔBSR-BRAF (151-766). All plasmids were cloned into pEGFPN3 plasmids linearized with HindIII and BamHI restriction enzymes using Gibson Assembly. A point mutation (R188L for BRAF, R89L for CRAF; denoted by an asterisk) that disrupts RAS binding was introduced to all RBD domains by site-directed mutagenesis. For the kinase dead mutant constructs, a point mutation in the kinase domain (D594G) was introduced by site-directed mutagenesis. All mammalian cell constructs were sequenced by whole plasmid sequencing.

### Bacterial protein purification

All RAF constructs were expressed in Rosetta (DE3) cells. A single colony was used to inoculate an overnight preculture, which was diluted into 1 L of lysogeny broth (LB) medium containing appropriate antibiotics at OD of 0.05 and grown at 37 °C with shaking. At OD□□□ ≈ 0.6, protein expression was induced with 30 μM (isopropyl β-D-1-thiogalactopyranoside) IPTG and continued for an additional 5 hours at 32 °C. Cells were harvested by centrifugation at 4 °C, flash-frozen in liquid nitrogen, and stored at –80 °C until purification. For purification, cell pellets were resuspended in a lysis buffer prepared with Buffer A (20 mM HEPES, pH 7.4, 500 mM NaCl, 1 mM TCEP, 10% glycerol) and supplemented with 15 μg·mL^-1^ DNase and 1 mM phenylmethylsulfonyl fluoride (PMSF). Cells were lysed through sonication, and the lysate was clarified by centrifugation and subsequently purified using fast protein liquid chromatography (FPLC) on an NGC system (Bio-Rad). A Strep-Tactin affinity column (Strep-TactinXT 4Flow, IBA Lifesciences) was used to perform an initial capture of the protein which was eluted with a gradient of buffer A supplemented with 50 mM biotin. Selected fractions were dialyzed in buffer A and incubated with TEV protease overnight at 4 °C. Subtractive rebinding of the cleaved tag and uncleaved protein were removed through the reintroduction of the sample onto a Strep-Tactin column. Size exclusion chromatography (Superdex 75 Increase HiScale, Cytiva) equilibrated with buffer A was used to further purify RAF. Selected fractions were analyzed through SDS-PAGE, combined as appropriate, aliquoted and snap frozen in liquid nitrogen, and kept at –80 °C.

### Supported lipid bilayer formation

SLBs were prepared as described elsewhere^37^. Briefly, 80% 1,2-dioleoyl-sn-glycero-3-phosphocholine (DOPC) and 20% 1,2-dioleoyl-sn-glycero-3-phospho-L-serine (DOPS) dissolved in chloroform were dried under vacuum for 20 min, followed by a stream of nitrogen gas for 30 min. Lipids were rehydrated in deionized water at 1 mg·mL^-1^. Lipid solution was sonicated to form small unilamellar vesicles (SUVs). Glass coverslips (D263 Schott glass, Ibidi) were sonicated in a 1:1 isopropanol/water mixture for 15 minutes, then incubated in 2% warm Hellmanex III for 30 minutes. The glass substrates were etched in piranha solution (3:1 H_2_SO_4_/H_2_O_2_) for 10 minutes. The coverslips were then assembled onto six-channel flow chambers (sticky-Slide VI 0.4, Ibidi). SUV solutions diluted at 0.25 mg·mL^-1^ in phosphate buffered saline (PBS, 10.1 mM of Na_2_HPO_4_, 1.76 mM KH_2_PO_4_, 137 mM NaCl, 2.68 mM KCl, pH 7.4.) was incubated with freshly etched coverslips for 45 min. Excess SUVs were rinsed with PBS, followed by buffer exchange with binding buffer (40 mM HEPES, 150mM NaCl, 5mM MgCl_2_).

### TIRF microscopy

Binding measurements on SLBs were performed on a Nikon Eclipse Ti-2 inverted microscope using an Apo 100X oil immersion lens. SLBs were illuminated in a total internal reflection mode with a 488 nm laser (Coherent) at 0.5 mW, passed through a Chroma ZET488/10x cleanup filter and directed with a Chroma zt488/640rpc dichroic mirror. Emitted fluorescent light passed through the same dichroic mirror and two stacked emission filters (ZET488/640m and ET535/70m, Chroma) and was detected by an EMCCD camera (iXon Life 897, Oxford Instruments).

### TIRF Binding measurements

Binding assays of mNG-fused RAF constructs on SLBs were performed in binding buffer (40 mM HEPES, 150mM NaCl, 5mM MgCl_2_, pH 7.4) supplemented with 0.1 mg·mL^-1^ casein at 23 °C. Binding kinetics were acquired with 10 second intervals. RAF was diluted based on the concentration of mature mNG, which was determined by a peak absorbance at 506 nm. Raw TIRF intensities were subtracted by the background intensity (the average intensity of the first three images before RAF injection). Intensity from solution-phase RAF was determined in separate experiments with isolated mNG and subtracted after maturation normalization. The TIRF intensities were finally converted to membrane-bound surface densities of RAF molecules by multiplying the conversion factor, determined by TIRF-FCS calibration curve (see Figure 1S). The equilibrium surface density was determined by fitting the binding kinetics to a two-species adsorption model. The partition coefficient, *K_p_*, is defined by the following equation^22^:

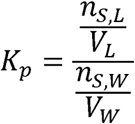

with *n_S,L_* and *n_S,W_* representing the number of moles of solute (RAF in this study) present in the lipid and aqueous phases, respectively. *V*_L_ and *V*_W_ denotes the volumes of the lipid and aqueous phase, respectively. *n*_S,L_ was calculated from the equilibrium surface density of RAF obtained from the binding kinetics. A lipid bilayer height of 4 nm was used to calculate the volume of the lipid phase. The bilayer surface area and flow cell volume were calculated based on the physical dimensions of the Ibidi flow chambers provided by the manufacturer.

### Fluorescence correlation spectroscopy

To determine mNeonGreen density from TIRF intensity, lipid-tethered His_10_-mNG, an mNG construct with an N-terminal affinity tag made of a sequence of 10 histidine residues to bind metal ions, was measured using both TIRF microscopy and FCS. SLBs comprised of 96% DOPC and 4% 1,2-dioleoyl-sn-glycero-3-[(N-(5-amino-1-carboxypentyl)iminodiacetic acid)succinyl] (nickel salt) (Ni-DOGS) were prepared in the same manner as other bilayers. Various surface densities of mNG on SLBs were obtained by incubating varying concentration of mNG in tris buffered saline (TBS, 20 mM Tris-HCl, 150 mM NaCl, pH 7.4) and washed with TBS. After acquiring TIRF intensity with the same imaging condition used in RAF binding assays, FCS measurements were immediately performed. A white light laser (NKT Photonics) was reflected from a dichroic mirror (zt488rdc, Chroma) and passed through a wavelength cleanup filter (LL01-488, Semrock) for wavelength selection. The laser was coupled to a single-mode polarization maintaining optical fiber (Thorlabs), collimated via a reflective collimator (RC08FC-P01, Thorlabs), and delivered to the back port of the microscopy. A Chroma ZT405/488/561/640rpc dichroic filter cube directed excitation and emitted light to and from the sample through the Apo 100X oil immersion TIRF lens. Emitted light is filtered through a 50 μm pinhole, collimated, split by a dichroic mirror (550lxpr, Chroma) and one last emission filter (ET520/40m, Chroma). The emission was focused for detection by an avalanche photodiode (SPCM-AQRH-16, Excelitas Technologies). FCS traces were recorded by a PicoHarp 300 TCSPC module. Data were converted in MATLAB (MathWorks) and fitted in Prism (GraphPad). Autocorrelation curves, *G*(*τ*) were adequately fitted with two-species diffusion to account for any population of less mobile long-residence particles arising from heterogeneity of His_10_ tagging chemistry:

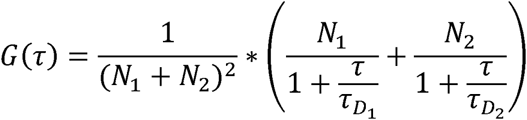

where *N*_1_ and *N*_2_ represents the average number of fluorescent molecules in fast and slow molecules, *N*, is the sum of *N*_1_ and *N*_2_. τ represents lag time, τ*_D_*_1_ represents characteristic diffusion within the two-dimensional focal spot, respectively. The average of total fluorescent diffusion time of fast species, τ*_D_*_2_ represents characteristic diffusion time of slow species. Diffusion coefficient for i^th^ species, *D_i_*, is calculated by the following equation:

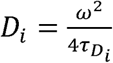

with *ω* representing the focal radius of the excitation laser.

### Live cell imaging

HEK293T cells were cultured in high glucose DMEM supplemented with 10% heat-inactivated FBS (HIFBS) and 1% penicillin/streptomycin at 37 °C under 5% CO2. For live cell imaging, HEK293T cells were plated onto 35 mm glass bottom dishes at a density of 1×10^6^ cells and incubated overnight at 37°C under 5% CO_2_. Cells were transfected with 1 µg of the indicated RAF plasmid supplemented with 7 µg of noncoding empty plasmid (pTT3) using calcium phosphate method. The following day, cells were washed twice with PBS and stained with CellMask™ Deep Red Plasma Membrane Stains (Invitrogen) for 5 minutes according to manufacturer protocols. For treatment with GDC-0879, cells were incubated with 2 µM of inhibitor for 2 hours before following staining protocol and sustained inhibitor concentration while imaging. Cells were kept in culturing media without phenol red and imaged using a 63x oil objective on an inverted Leica Stellaris 5 inverted confocal microscope. Images were acquired using HyD detectors with the following excitation/emission settings for each fluorescence channel: mNG-504 nm /509-622 nm, CellMask™ Deep Red Plasma Membrane Stain603 nm/638-750 nm. To quantify plasma membrane enrichment, we manually drew multiple segmented lines from extracellular to cytosolic space while crossing the plasma membrane. Plasma membrane regions at the junctions between transfected cells were excluded to avoid double counting. Fluorescence intensity profiles of the red channel (plasma membrane) were extracted using ImageJ. To ensure consistent measurement paths, the same lines were applied to the green channel (RAF-mNG), and fluorescence intensities were extracted as above. Background subtraction was performed by applying the same protocol to untransfected cells. Using custom MATLAB scripts, we identified the peak fluorescence intensity points corresponding to the plasma membrane in the red channel and mapped these positions onto the green channel data. Plasma membrane enrichment was calculated by dividing the average background-corrected fluorescence intensity of RAF-mNG at the plasma membrane by the average cytosolic RAF fluorescence intensity. Measurements were performed on 20 cells per condition.

## Supplementary Material Description

File name: Supporting Information.docx

The supporting information provides data on TIRF-FCS calibration experiments, protein purification of CRAF constructs, electrostatic properties of BSR and CSR, electrostatic surface representation of BSR-RBD-CRD and CSR-RBD-CRD, net charge changes with progressive N-terminal truncation of BSR, and confocal microscopy images showing subcellular localization of BRAF constructs in HEK293T cells after treatment of GDC-0879.

## Author Contributions

**Vasili Revazishvili:** investigation, writing - original draft, validation, methodology, formal analysis, data curation, conceptualization, resources. **Alexia Morales:** investigation, writing - original draft, methodology, validation, visualization, formal analysis, data curation, conceptualization, resources. **Bryn Baxter:** investigation, methodology, formal analysis, resources. **Julian Grim:** investigation, methodology, validation, formal analysis, data curation, resources. **Ani Chakhrakia:** investigation, conceptualization, visualization, validation, formal analysis, data curation, resources. **Andres Jimenez Salinas:** investigation; methodology, validation, resources. **Angelica M. Riestra:** investigation, methodology, resources, funding acquisition. **Young Kwang Lee:** Supervision, project administrator, writing-original draft, investigation, formal analysis, conceptualization, funding acquisition. All authors provide inputs on the manuscript writing.

## Acknowledgments

This study was supported by NSF CAREER Award MCB-2145852 to Y.L and by a Research Project Grant to A.M.R from the National Institute on Minority Health and Health Disparities of the National Institutes of Health under Award Numbers S21MD010690 (SDSU HealthLINK Endowment) and U54MD012397 (SDSU HealthLINK Center). The content is solely the responsibility of the authors and does not necessarily represent the official views of the National Institutes of Health. A.M.R was also supported by a Prebys Research Heroes Grant from the Prebys Foundation A.J.S. was supported in part as a Fellow of the Rees-Stealy Research Foundation. B.B. was supported in part by an Association for Women in Science-San Diego Scholarship.

## Data Availability Statement

The data that support the findings of this study are available from the corresponding author upon reasonable request.

