## Supporting Information for "Quantitative Membrane Binding Assays Reveal an Inhibitory Role for the BRAF-Specific Region in CRD and Lipid Interaction"

**Supplementary Figures and Tables**

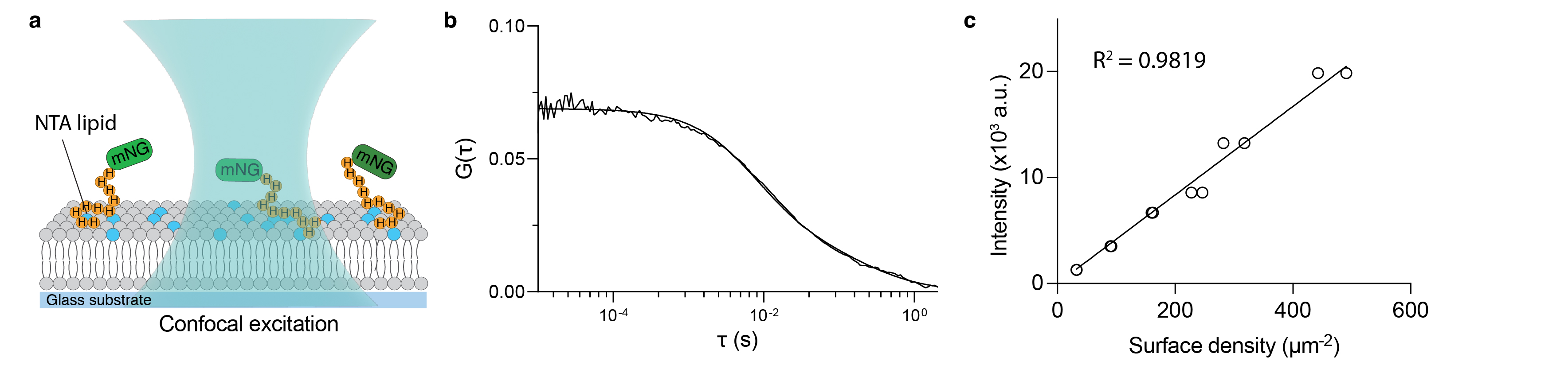

**Figure S1.** Fluorescence correlation spectroscopy (FCS)-based calibration of RAF surface density. (a) Schematic of the FCS setup showing mNeonGreen (mNG) proteins tethered to DGS-NTA(Ni) lipids via an N-terminal His_10_ tag. As mNG diffuses through the illuminated confocal volume, fluctuations in fluorescence intensity are recorded and used to calculate the autocorrelation function G(τ), enabling quantification of diffusion dynamics and surface density. (b) Representative autocorrelation curve and corresponding fit for mNG tethered to supported lipid membranes in TBS buffer (20 mM Tris, 150 mM NaCl, pH 7.4) supplemented with 0.1 mg/mL casein. Due to the heterogeneous nature of the His_10_ tag, at least two distinct diffusive populations were observed. The data were best fit using a two-species diffusion model for a two-dimensional system, yielding diffusion coefficients *D*_1_= 0.90 and *D*_2_= 0.03 μm²/s and average particle counts *N*_1_ and *N*_2_ of 11.0 and 3.5 for each population respectively. The total number of particles, *N*, is sum of *N*_1_ and *N*_2_. The surface density was calculated by dividing *N* by the area of two-dimensional focal area. (c) Calibration curve showing a linear relationship between FCS-derived surface density and total internal reflection fluorescence (TIRF) intensity for mNG-functionalized supported membranes.

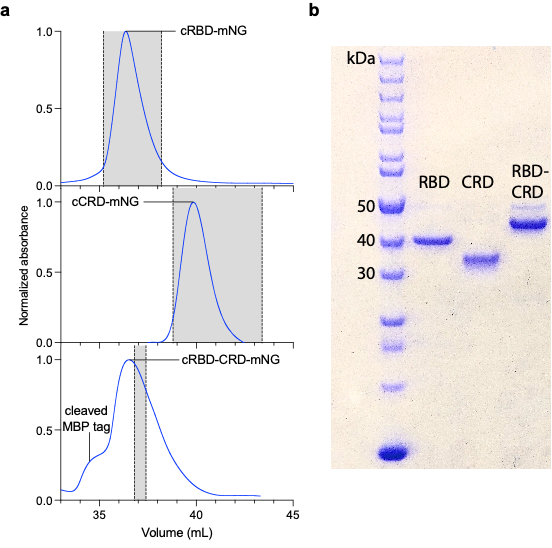

**Figure S2.** CRAF constructs were robustly expressed, soluble, and highly pure. (a) Size exclusion chromatography (SEC) elution profiles of CRAF-mNG constructs monitored by 280 nm absorbance. Grey shaded areas indicate collected fractions. For cRBD-CRD-mNG, the cleaved MBP tag eluted prior to the protein of interest. Only fractions enriched in the target protein were collected, as confirmed by SDS-PAGE. The expected elution volumes were 37 mL for cRBD–mNG, 39 mL for cCRD–mNG, and 36 mL for cRBD-CRD-mNG. All constructs eluted near these predicted volumes. (b) SDS-PAGE analysis showed discrete bands at expected molecular weights. The RBD and CRD constructs were virtually pure. The cRBD-CRD-mNG sample exhibited a minor band at ~50 kDa, corresponding to the cleaved MBP tag. Estimated purity of cRBD-CRD-mNG was ~90% based on band intensity. Note that proteins were diluted for binding assays based on the concentration of matured mNG, and non-fluorescent protein impurities do not contribute to fluorescence signal or influence dilution accuracy.

**Table S1.** Electrostatic properties of BSR and CSR.

|  | pI^1^ | Net charge at pH 7.4^1^ | No. of negatively  charged residues | No. of positively  charged residues |
| --- | --- | --- | --- | --- |
| BSR | 4.13 | -13.1 | 20 | 9 |
| CSR | 5.28 | -1.5 | 5 | 6 |

^1^ Electrostatic parameters were calculated using the Isoelectric Point Calculator (IPC 2.0; <https://isoelectricpoint.org>).

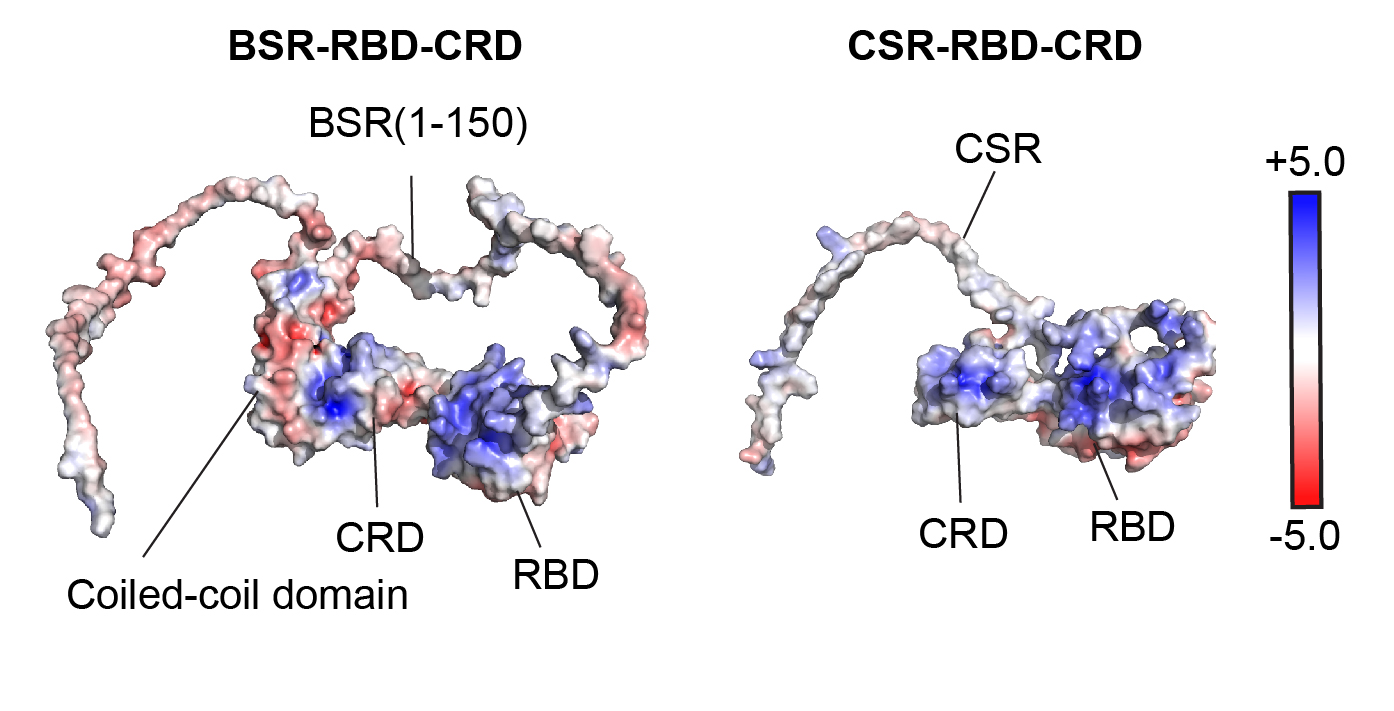

**Figure S3.** Electrostatic surface potential of BSR-RBD-CRD and CSR-RBD-CRD mapped onto the Alphafold2 structures. The BSR contains multiple negatively charged residues distributed across its sequence, whereas the CSR displays a relatively neutral electrostatic surface potential.

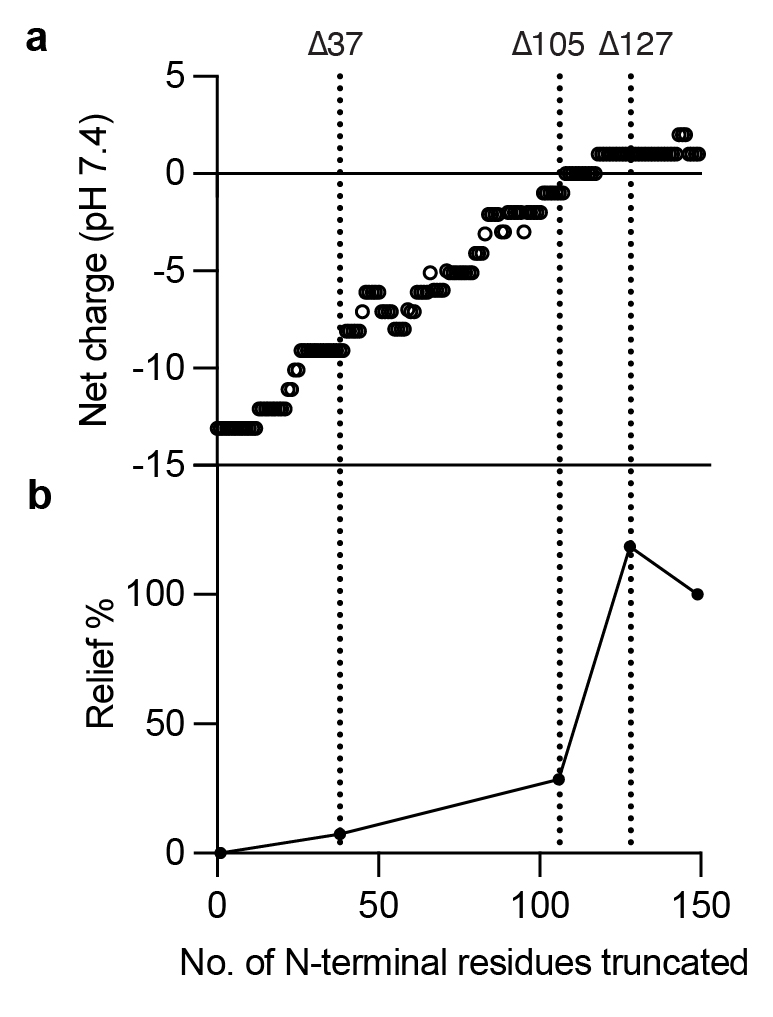

**Figure S4.** (a) Net charge of the isolated BSR and (b) relief of lipid interaction in BSR-RBD-CRD constructs with successive N-terminal truncation at single–amino acid resolution. A sharp relief is observed upon truncation of the first 127 residues, coinciding with a shift in net charge from negative to positive.

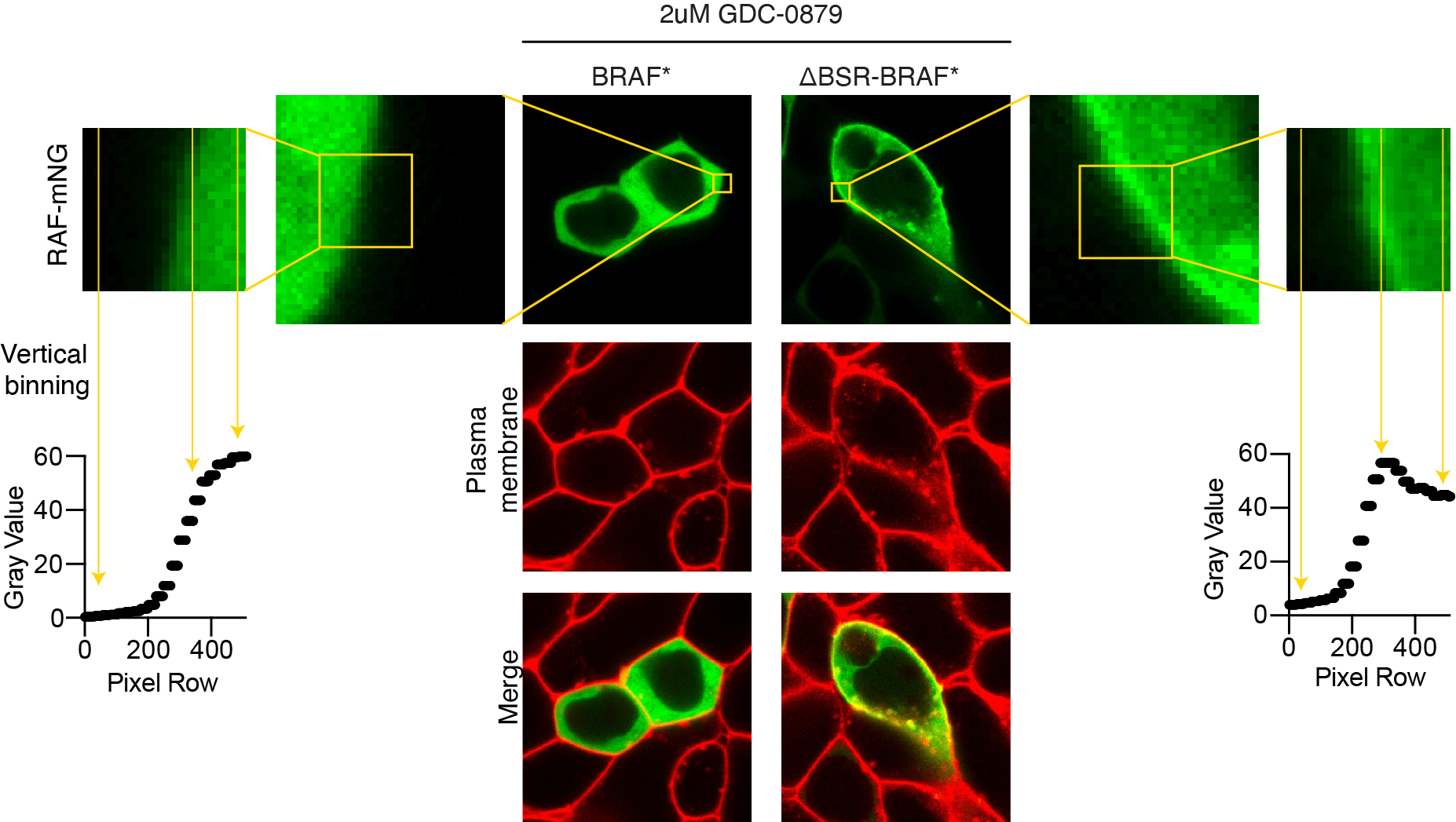

**Figure S5.** The BSR suppresses CRD–lipid interaction when CRD autoinhibition is weakened by RAF inhibitor treatment. Representative confocal microscopy images of HEK293T cells transfected with either FL-BRAF* or ΔBSR-BRAF*. Cells were treated with 2 μM GDC-0879 for 2 hours. The green channel shows mNeonGreen-tagged BRAF constructs, and the red channel indicates plasma membrane staining with CellMask Deep Red. A subpopulation of ΔBSR-BRAF*-expressing cells exhibits plasma membrane enrichment, as highlighted in the zoomed-in panel (top right). The inset graphs represent a vertically binned one-dimensional horizontal intensity profile of the zoomed-in panels. Vertical binning was performed by calculating the average pixel intensities along each column of the image.
